# Development of a noninvasive olfactory stimulation fMRI system in marmosets

**DOI:** 10.1101/2024.04.21.590223

**Authors:** Terumi Yurimoto, Fumiko Seki, Akihiro Yamada, Junnosuke Okajima, Tomoyuki Yambe, Yoshiaki Takewa, Michiko Kamioka, Takashi Inoue, Yusuke Inoue, Erika Sasaki

## Abstract

Olfactory dysfunction is associated with aging and the earliest stages of neurodegenerative diseases, such as Alzheimer’s and Parkinson’s diseases; it is thought to be an early biomarker of cognitive decline. In marmosets, a small non-human primate model used in brain research, olfactory pathway activity during olfactory stimulation has not been well studied because of the difficulty in clearly switching olfactory stimuli inside a narrow MRI. Here, we developed an olfactory-stimulated fMRI system using a small-aperture MRI machine.

The olfactory presentation system consisted of two tubes, one for supply and one for suction of olfactory stimulants and a balloon valve. A balloon valve installed in the air supply tube controlled the presentation of the olfactory stimulant, which enabled sharp olfactory stimulation within MRI, such as 30 seconds of stimulation repeated five times at five-minute intervals. The olfactory stimulation system was validated in vivo and in a simulated system. fMRI analysis showed a rapid increase in signal values within 30 s of olfactory stimulation in eight regions related to the sense of smell. As these regions include those associated with Alzheimer’s and Parkinson’s diseases, olfactory stimulation fMRI may be useful in clarifying the relationship between olfactory dysfunction and dementia in non-human primates.

## Introduction

Olfactory dysfunction is associated with neurodegenerative diseases such as Alzheimer’s disease (AD) and Parkinson’s disease (PD)^1-3^, and has been reported in 96% of patients with PD and 90% of patients with AD^3^. Moreover, the dysfunction is very common in the older population, being present in >50% of individuals aged between 65 and 80 years and in 62–80% of those >80 years of age^4^. Olfactory dysfunction precedes the onset of dementia in both AD and PD^3^, and it has been reported that olfactory function would be an important sensory biomarker of cognitive change in longitudinally followed older adults (aged 75 years at the start of the study)^1,5^. An olfactory experimental platform is indispensable to reveal the mechanisms underlying the association between olfactory dysfunction and brain disorders.

The common marmoset (*Callithrix jacchus*) has a relatively short lifespan among primates, and its small size (300–500 g) and fecundity make it an increasingly interesting human aging model^6^. Marmosets have a well-developed olfactory system for various behaviors, including reproduction, communication, and food selection^7^. Marmosets, like rodents, have an accessory olfactory system, but the accessory olfactory system is not directly relevant in assessing olfactory function^7,8^. This indicates that the anatomical and functional differences in olfaction between humans and marmosets are negligible and that there are few obstacles to using marmosets for olfactory research^7^. Marmosets have many human-like characteristics in various aspects, such as an enlarged temporal lobe, a highly developed prefrontal cortex, hierarchically organized sensory areas^9^, and other brain structures typical of primates, as well as a rich social repertoire and cognitive behaviors similar to those of humans^10,11^. Together, these data indicate that marmosets are excellent candidates for studying the relationship between olfaction and brain function by functional MRI (fMRI), which is a powerful tool in neuroscience and cognitive function research and is used in basic, translational, and clinical research^12-14^.

Anatomical olfactory systems have been extensively studied in various species, including nematodes, drosophila, zebrafish, mice, rats, dogs, and humans^3,15-19^. In mammals, the specific function, organization, and genetic complexity of the olfactory system vary widely among species^3^. For example, mice have about 1130 olfactory receptor (OR) genes, rats have 1207 OR genes, and dogs have 811 OR genes, while humans have 396 OR genes^20^. Non-human primates are similar to humans, with chimpanzees having 380 OR genes and marmosets having 366 OR genes^20^. There are anatomical differences between human and rodent olfactory systems, such as a higher ratio of olfactory bulb volume to brain volume in rodents than in humans^3^. Although olfactory nerve projection patterns are preserved across species, the anatomical shape of the nasal cavity and the distribution of olfactory nerves differ between rodents and humans^21^. However, the similarity in the number of OR genes and the nasal anatomy between marmosets, which are non-human primates, and humans suggests that marmosets may be a useful animal model for olfactory research^21,22^. Moreover, it has been reported that mice and rats are more resistant to age-related changes in the olfactory system^23,24^. On the contrary, olfactory discrimination deficits can be experimentally induced by the toxin MPTP (1 □methyl □4 □phenyl □1,2,3,6 □tetrahydropyridine) in marmoset models of PD^25^. Marmosets have also emerged as a variety of disease models using genetically modified animal creation techniques^26-28^. Therefore, marmosets are excellent candidates for studying age-related changes in olfaction.

Marmosets can obtain superior signal-to-noise ratio and resolution images compared with macaques, as marmosets can be applied to ultra-high field MRI for rodents. However, the small size of the MRI machine limits the use of equipment for stimulation and observation^29^. Therefore, previous fMRI studies on marmosets were accomplished by producing special devices^29-31^. However, the application of these special devices across all experimental conditions is difficult due to limitations such as the need for anesthesia, invasive head immobilization, and the caliber of the MRI equipment used. Recently, Seki et al. created a non-anesthetic, noninvasive holder that can perform awake fMRI in small-bore diameter MRI^32^, making fMRI studies feasible for a wider range of purposes. There are also difficulties in developing olfactory stimulant delivery devices that can tightly control the delivery and removal of olfactory stimulants. In particular, non-human primate faces do not have nose corns, and developing a device that does not leak stimulants is difficult. Previously, an olfactory stimulation fMRI study in marmosets was conducted by Ferris et al., in which male marmosets sniffed urine associated with female sexual cycles and sex hormones, and fMRI was used to investigate brain activity related to emotions in the thalamus and other areas^31,33^. However, because prolonged stimulation induces olfactory adaptation^34,35^, olfactory stimulation with a smell-impregnated cloth placed in front of the marmoset’s face for 7 min may be unsuitable for studies interested in the brain activity of the main olfactory pathway. Furthermore, in previous studies, the animals were anesthetized to restrain animals securely prior to the acquisition of imaging^31,33^, and the effect of anesthetics on brain activity cannot be ruled out even when performing fMRI after recovery. To avoid the effects of olfactory adaptation, our device was designed to analyze brain activity for short periods and reset olfactory stimuli. This allowed us to increase the number of trials within a limited time period, during which the animals could be held without anesthesia. In this study, to achieve fMRI with olfactory stimulation in a completely noninvasive awake marmoset without anesthesia, we improved the device to allow for olfactory stimulation that can be inside MRI and then performed controlled olfactory stimulated fMRI.

## Results

### Animal holder

The design of the animal holder was improved from a previous report using 3D CAD (Fusion 360; Autodesk, San Francisco, CA)^32^. The helmet was divided into two parts: inner and outer. The inner helmet had a built-in receiver coil, which improved the signal-to-noise ratio duo to reduce the gap between the head and the coil (Takashima Seisakusho Ltd., Tokyo, Japan). The gap between the helmets was adjusted for each marmoset by inserting sponges between the helmets. Marmosets were placed in the sphinx position. A schematic representation of the olfactory stimulation circuit is shown in Fig. 1A. Two holes were made on both sides of the chin rest, and the tubes were threaded through them. The tube tips were designed to be slightly bent for placement near the marmoset’s nose (Fig. 1B, C). One tube was used as the supply tube and the other as a suction tube. A balloon valve was placed 50 mm away from the tip of the supply tube. The supply tube was connected to an olfactory test unit (O’ Hara & Co., Ltd., Tokyo, Japan), and stimulation was supplied by a compressor.

**Figure 1.**
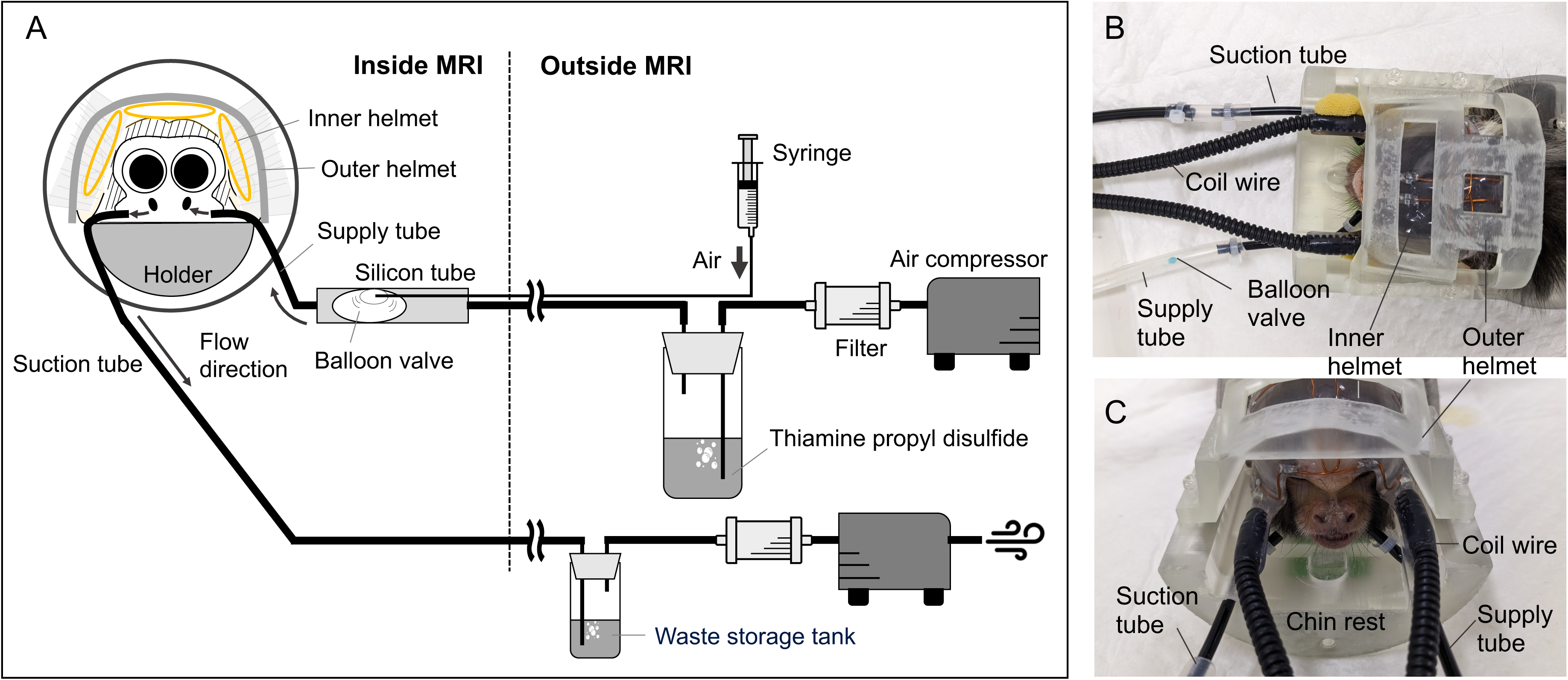
3D printed holder for MRI and olfactory stimulation circuit. The left side of the animal view shows the supply tube, and the right side shows the suction tube. (A) Schematic diagram, (B) top view, and (C) front view.

### Visualization of stimulus airflow

The experimental environment for the in vivo experiment was constructed using the designed holder restrainer and animal mock-ups (Fig. 2A–C). The results shown in Fig 2D and 2E are the airflow images for each second with and without suction. Figure 2 (D and E) shows the results of an experiment to visualize the stimulation flow in an MRI-simulated environment to evaluate the utility of the suction tube. Figure 2 (D1–D3) shows continuous images of the gas flow for approximately 1 second of visualizing the stimulation flow, only air delivery and no suction. The particles did not vent, became a swirling flow, and stagnated around the marmoset’s face. The results shown in Fig. 2E1–E3 are 1-s sequences of airflow with suction. The outgoing smoke passed in front of the marmoset’s nose and was smoothly exhausted in a laminar flow. No particles were observed to remain in the air for long periods.

**Figure 2.**
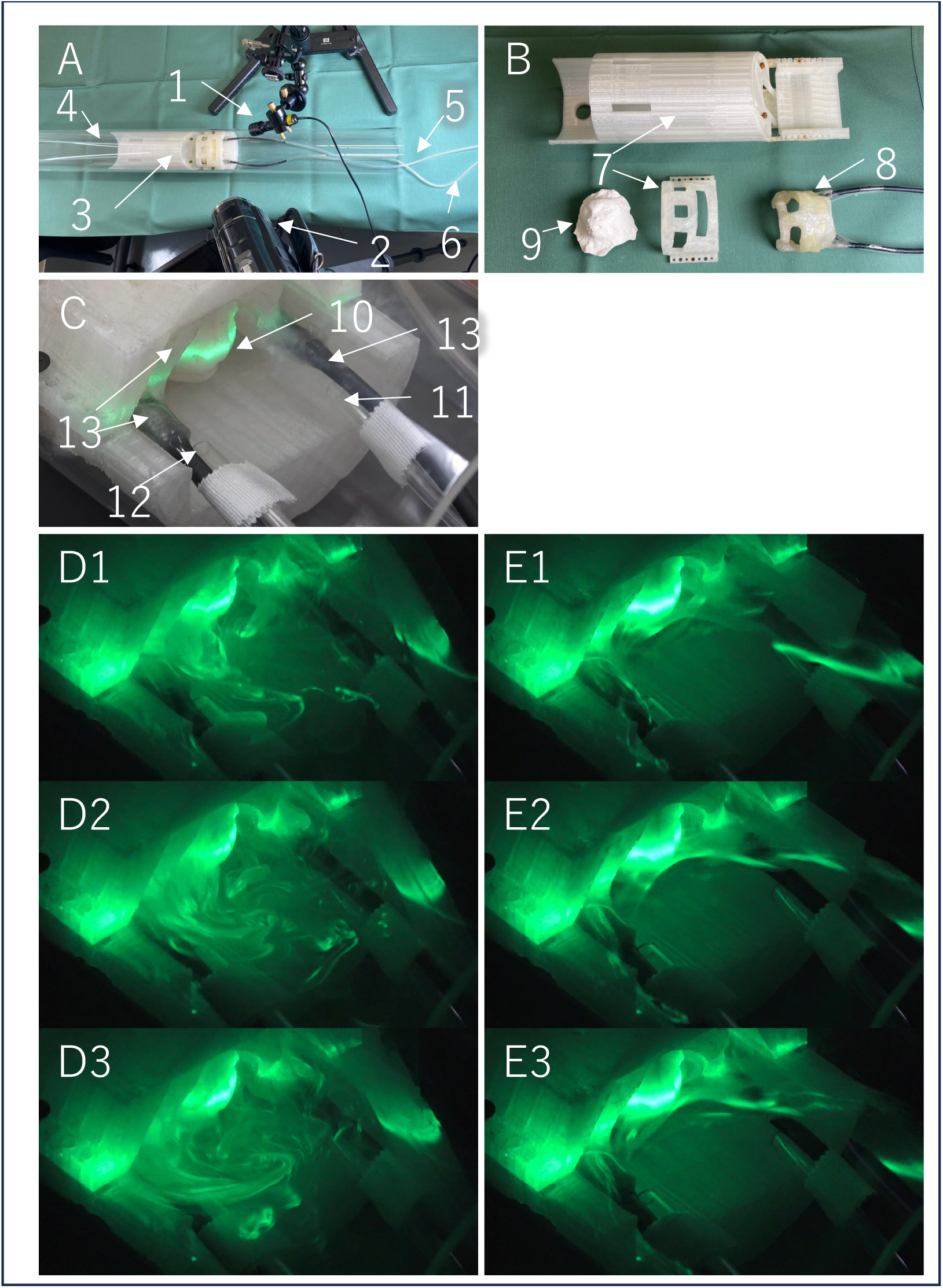
Flow visualization experiment to evaluate the usefulness of suction. A: Overall view of the simulated MRI environment. laser light source (1), video camera (2), marmoset holder (3) in a simulated MRI environment (4), supply tube (5), and suction tube (6). B Marmoset Holder Components. Marmoset holder and outer helmet (7) and inner helmet (8) with a marmoset face model (9). C: Enlarged view of the face. The laser sheet was placed horizontally at the marmoset nose level (10), and the supply tube (11) and suction tube (12) were placed parallel to the coil wire (13). D: Continuous images showing the gas flow for 1 s when only air was supplied. Odors stagnate and flow in circles. E: 1-s continuous images of the air delivery and suction. The delivered odor flowed smoothly through the nose of the marmoset in laminar flow.

### Motion evaluation

Movement of the head position coordinates from the start of each fMRI scan was observed. The movements of a representative animal, I817, the slightest movement among the trained marmosets, of the six extracted motion parameters (x, y, z translation, and rotation) during the runs are shown in Fig. 3. The marmoset kept the movements less than 0.5 mm in all trials. Although head movements increased slightly when the animals began stimulation (30 s), the movements were within the acceptable range, and a stable restraint was observed. The movements in the anterior-posterior direction were the largest, whereas those in the vertical and lateral directions were approximately 0.1–0.2 mm. Other marmosets’ movements were less than 0.5 mm, and even the trial with the most movement was 0.3 mm (Supplemental Fig. 1).

**Figure 3.**
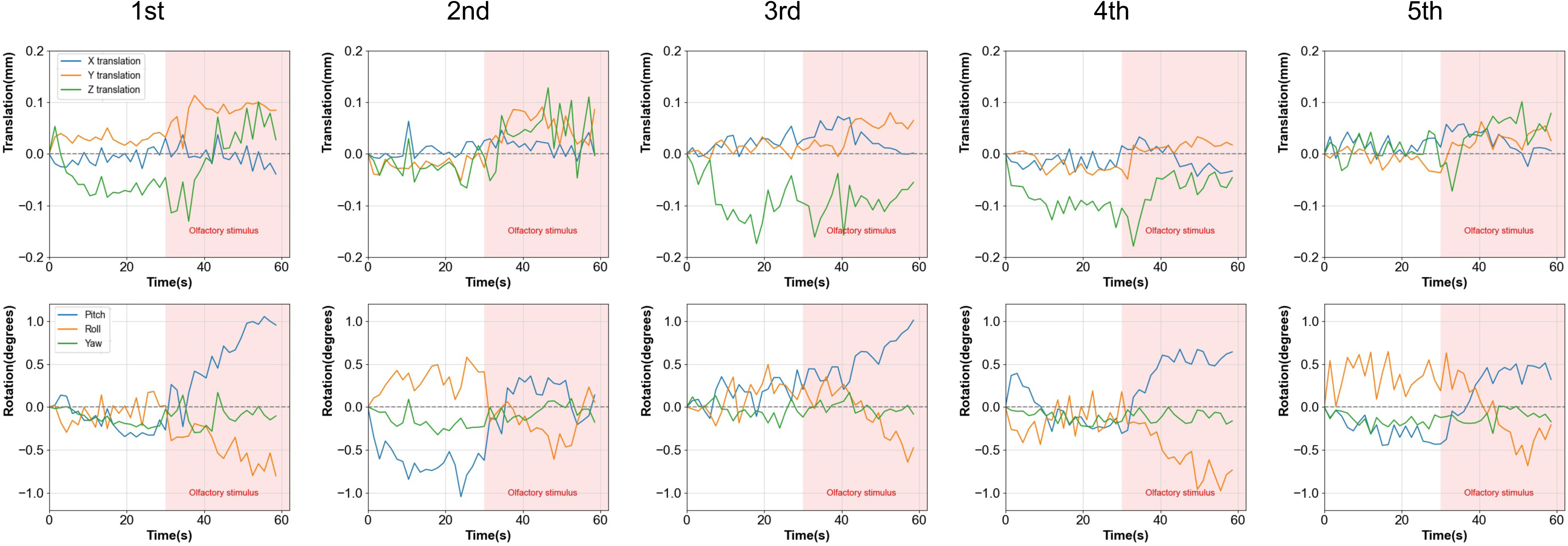
Estimates of head motion during MRI imaging of a marmoset. Estimates of parallel translation (top row) and rotation (bottom row) are shown, and each column shows the results of 40 individual-volume fMRI runs. All images were captured in five trials on the same day. Functional imaging was performed in five runs for each animal (the interval between each run was 5 min). The red background area indicates the time of olfactory stimulation. All the data were acquired from the I817.

### Olfactory fMRI

The BOLD signal value of the olfactory bulb (OB) was immediately increased by olfactory stimulation, followed by a decrease in the OB’s response when the stimulation was stopped in 7-min scan (Fig. 4A). Anterior olfactory nucleus (AON) and both side orbitofrontal (Orbi) also showed a similar trend (Fig. 4B, H, I). The BOLD signal decreased from 90 s after the stimulation stopped and was at its lowest value at approximately 240 s after the stimulus. However, piriform (Piri), anterior cortical amygdaloid nucleus (ACo), mediodorsal nucleus of thalamus (MD), and both side entorhinal (Eth) did not exhibit this trend (Fig. 4C, D, E, F, and G). The BOLD signals of primary somatosensory (S1), primary visual cortex (V1), and auditory cortex (Aud) were independent of olfactory stimulation (Supplemental Fig. 2A, C, D). Gustatory cortex (Gu) showed a trend similar to that of OB (Supplemental Fig. 2B).

**Figure 4.**
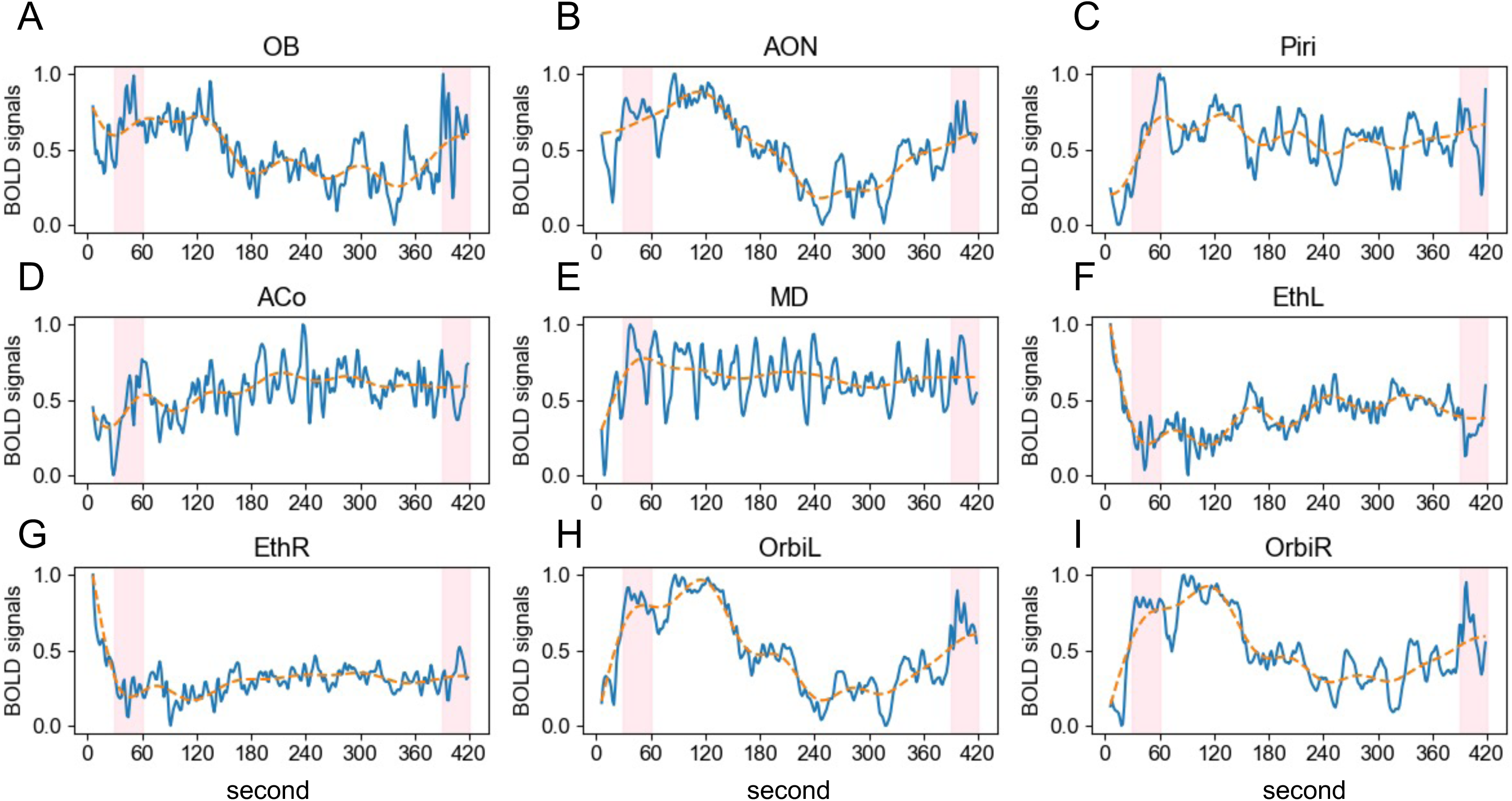
BOLD signals at the brain region with on/off olfactory stimulation. The blue line shows the normalized BOLD signal values acquired continuously for 7 min in the ROI. The orange dotted line represents the moving average. The pink areas indicate the times when the olfactory stimuli were administered. Olfactory stimulation was performed for 30 s, with stimulation intervals of 5 min 30 s. (A) OB: olfactory bulb, (B) AON: Anterior olfactory nucleus, (C) Piri: piriform cortex, (D) ACo: anterior cortical amygdaloid nucleus, (E) MD: mediodorsal nucleus of thalamus, (F) EthL: left entorhinal cortex, (G) EthR: right entorhinal cortex, (H) OrbiL: left orbitofrontal cortex, (H) OrbiR: right orbitofrontal cortex

Figure 5A-C shows the normalized BOLD signals of the 13 brain regions for each marmoset and figure 5D shows three marmosets average. Each marmoset showed a mild baseline increase before stimulation, which was stable and slight. Several regions of interest (ROI) signals increased after the stimulation. In particular, AON and right orbitofrontal (OrbiR) showed a large signal increase in all three marmosets and Piri in two. On the contrary, the increase was less than 1% for ACo, left orbitofrontal (OrbiL), and V1 at I7000; OB, Piri, and Gu at I814; and OrbiL and V1 at I817. The ROI with the lowest rate of increase was V1 followed by Orbi L. There were significant differences in the AUC, pre- and post-stimulation in the OB, Piri, ACo, right entorhinal (EthR), and OrbiL (*p*<0.05), and AON, left entorhinal (EthL), and OrbiR (*p*<0.01) (Fig. 6). No significant differences were found in MD, S1, Gu, or Aud. The olfactory pathways and responses in each region are shown (Fig. 7A). Significant differences were observed in all regions except the MD of the olfactory pathway, whereas no significant differences were found in regions unrelated to the olfactory pathway. Despite being the endpoint of the olfactory pathway, the Orbi response rate was faster than that in the other olfactory regions (Fig. 7B). In each run, the unresponsive trials did not always appear in the final part of the trial.

**Figure 5.**
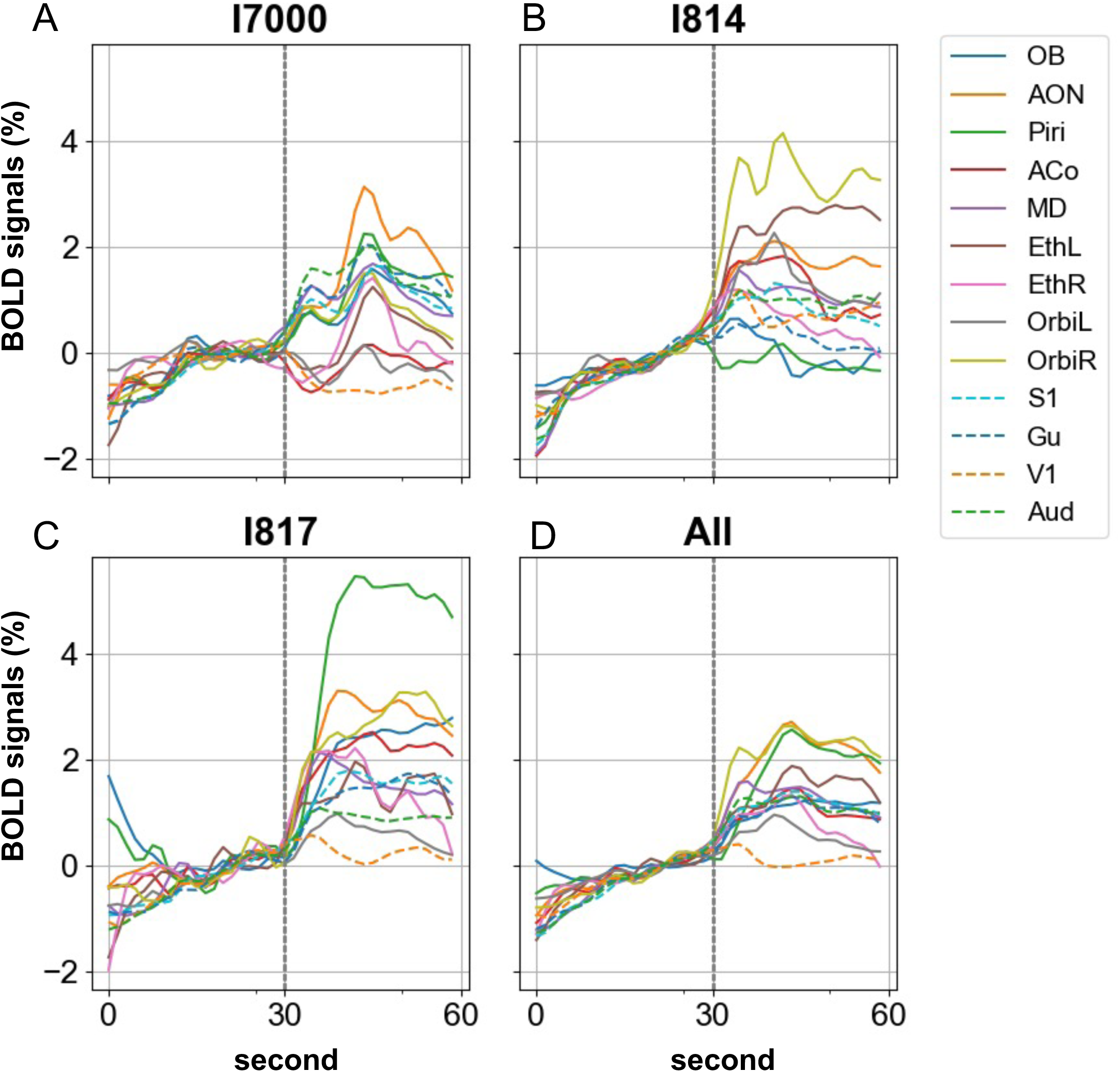
BOLD signals obtained from three marmosets. Three marmosets (A-C) and average (D) signals were shown. The dashed line at 30 s is the start time of olfactory stimulation. The curves represent averages of the signals in each region. OB: olfactory bulb, AON: Anterior olfactory nucleus, Piri: piriform cortex, ACo: anterior cortical amygdaloid nucleus, MD: mediodorsal nucleus of thalamus, Eth: entorhinal cortex, Orbi: orbitofrontal cortex, S1: primary somatosensory cortex, Gu: gustatory cortex, V1: primary visual cortex, Aud: auditory cortex

**Figure 6.**
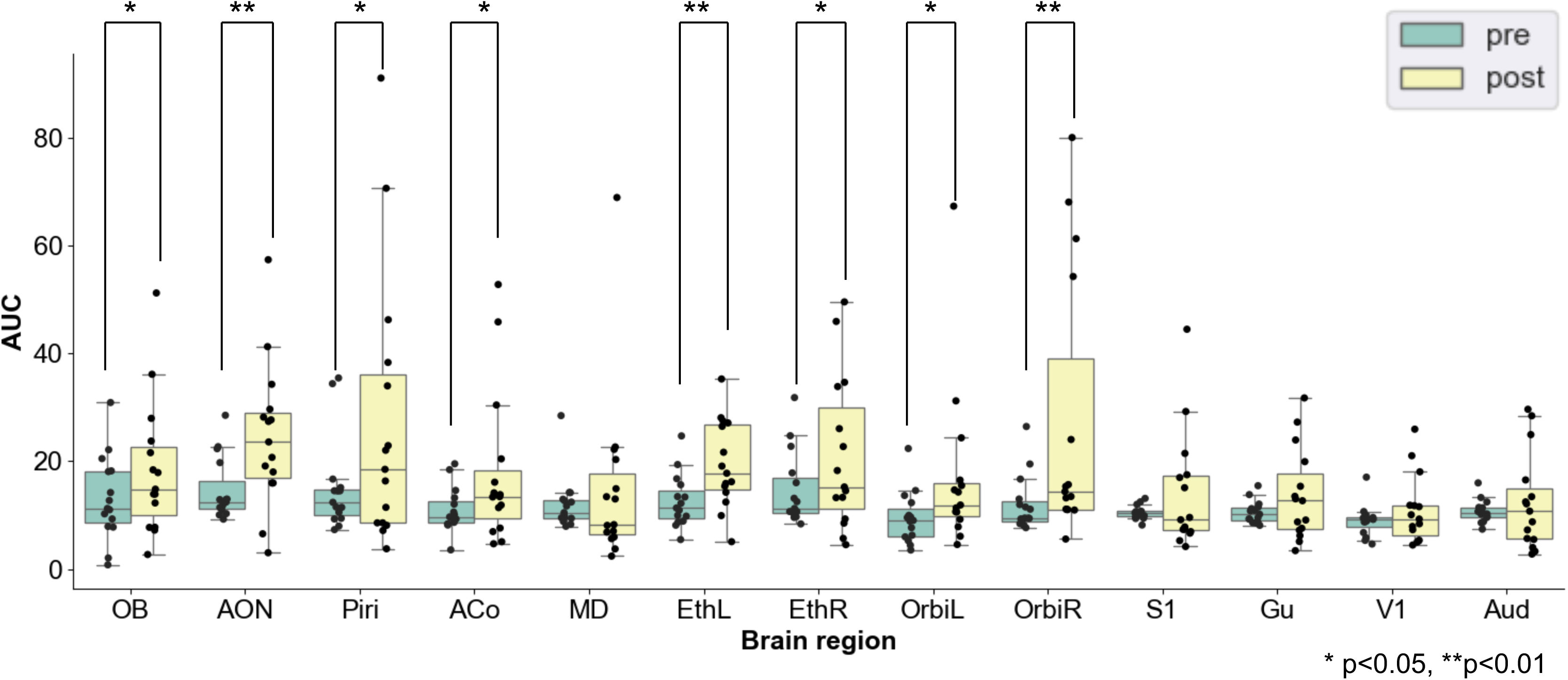
Box-and-whisker plot of AUC (Area Under Curve) before and after stimulation in each region. OB: olfactory bulb, AON: Anterior olfactory nucleus, Piri: piriform cortex, ACo: anterior cortical amygdaloid nucleus, MD: mediodorsal nucleus of thalamus, Eth: entorhinal cortex, Orbi: orbitofrontal cortex, S1: primary somatosensory cortex, Gu: gustatory cortex, V1: primary visual cortex, Aud: auditory cortex, *P<0.05, **P<0.01

**Figure 7.**
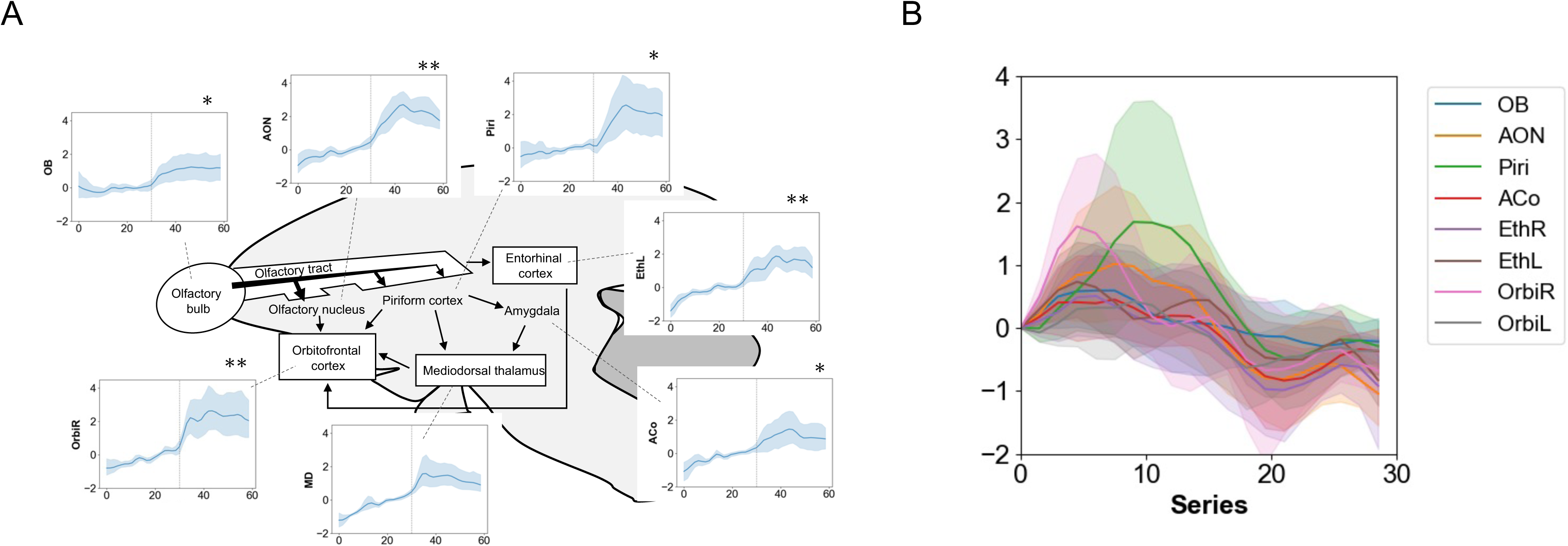
Olfactory pathway and brain reaction time after olfactory stimulation. (A) Schematic of the olfactory nerve pathways, asterisk marked areas with significant differences between pre- and post-stimulation. (B) Temporal changes in brain activity: The time from the start of the olfactory stimulation to the peak of the signal value. OB: olfactory bulb, AON: Anterior olfactory nucleus, Piri: piriform cortex, ACo: anterior cortical amygdaloid nucleus, MD: mediodorsal nucleus of thalamus, Eth: entorhinal cortex, Orbi: orbitofrontal cortex

## Discussion

In this study, we modified an existing fMRI holder for non-human primates to create a device capable of repeated olfactory stimulation in a short period and evaluated its performance in a mock circuit and in vivo. As olfactory adaptation is known to occur in a short period^34^, equipment that can swiftly control the olfactory stimulant supply is required to obtain olfactory fMRI data. In the first pilot study, a transvenous olfactory stimulation test^36^ was performed in marmosets; however, the activation of nerves in the olfactory network, including the olfactory bulb, was mild and did not provide clear results (Supplemental Figure 3). In the second pilot study, when olfactory stimulation was controlled without a balloon valve, even at a flow rate of 3 L/min (maximum flow), it took more than 1 min for us to detect the stimulation and 30 s for the stimulation to disappear. These pre-studies were unsuitable for analyzing brain function after short-term stimulation because of slow olfactory stimulation. Therefore, a specific device was needed to control the sharp stimulation. Attaching the balloon just before the air supply tube made it possible to turn the stimulation on and off sharply, and the stimulation was provided directly in front of the animal’s face.

To minimize the head motion of awake marmosets during ultra-high-field fMRI and improve spatiotemporal resolution, we designed a double-helmet holder with a helmet-integrated coil. The integrated coil-and-helmet design also made it possible to place the coil close to the brain, and the position between the coil and marmoset brain was stable. It allowed acquiring images with a 50% spatial signal-to-noise ratio improvement compared with images without the integrated coil, according to the data obtained in the preliminary study. From the results of five fMRI scans, all three marmosets demonstrated that the holder could minimize head movement together with sufficient pre-training, which led to minimal data being discarded owing to animal movement. Although there were individual differences between the animals, the animals with the supply tube near the nose showed slight head movements owing to the stimulant’s air pressure, sometimes hitting the animal’s whiskers or mouth. However, no significant changes were observed in the somatosensory cortex. Additionally, the spatial resolution for this study was 0.6 mm, and all trials had head movements of 0.3 mm or less below the resolution. Because the gas flow did not affect the body’s movement and did not affect brain functions except for the sense of smell., this device was suitable for olfactory stimulation. Another advantage of this device is that it can be used in experiments without unnecessary excitation or visual stimuli. When changing the presented olfactory stimulation, it is unnecessary to change the olfactory stimulant source in front of the animal, as in previous studies^31,33^.

To visualize the amount of residual gas in a physical environment equivalent to that in an MRI experiment, the gas was quickly removed from the vicinity of the stimulation target by suction, demonstrating the usefulness of aspiration. However, when the airflow rate was increased to the same level as in the actual animal experiment, the tracer particles diffused rapidly, making it difficult to visualize them. Therefore, in this model experimental system, we focused on the visualization of airflow with and without an exhaust route by lowering the flow rate from that in the animal experiment and conducted measurements under the experimental conditions that enabled visualization. The concentrations of the tracer particles required for visualization and the substances required for olfactory stimulation were different; therefore, it was unnecessary to match them in this experiment to demonstrate the usefulness of exhaust air. The inner diameters of the air supply and suction tubes used in the animal experiments were 2 mm, whereas the inner diameter of the tube used in the visualization experiments was 4 mm. The smoke for visualization tended to stick to the inner wall of the tubes, and when tubes with an inner diameter of 2 mm were used as air delivery tubes, it was not possible to send enough smoke for visualization. Olfactory stimulants from the supply tube passed through the marmoset’s nasal tip and were discharged from the suction tube in a laminar flow. If there was no suction and the olfactory stimulant was simply released into the marmoset’s nose, the olfactory stimulant would remain in the marmoset’s nose for an unintentionally long period of time, stimulating the sense of smell and slowing the response time of the experimental system. This indicates that suction is useful for this experimental system.

The signal increased in eight of the nine brain regions associated with the olfactory pathway after olfactory stimulation (Fig. 6). However, there was no significant increase in brain activity in regions that were not associated with olfactory functions, such as the somatosensory and gustatory cortices. Therefore, the purpose of this device, which is to specifically stimulate and assess olfactory function, was achieved. Furthermore, this device has two advantages: a high temporal resolution and ventilation around the marmoset is completed in each trial. In fact, the signals showed a significant difference within 30 s after stimulation, allowing evaluation with a high temporal resolution compared to previous studies^31,37^. Although it is known that prolonged continuous olfactory stimulation causes a loss of responsiveness in the Piri and AON^34,38^, the first half of the olfactory pathway, including the AON, was reactivated by repeated stimulation in this study. Thus, the 5-min interval time was sufficient, and the air around the marmoset was considered ventilated. During 7 min of continuous fMRI imaging, the BOLD signal dropped once and rose again when the second olfactory stimulus was provided in the four regions. In vivo studies demonstrated the effectiveness of this olfactory stimulator.

By contrast, the second stimulus to Piri did not sufficiently increase the raw value of the BOLD signal. In rats, the second and subsequent stimuli are weaker than the first, and applying the same stimulus several more times changes the signal in a negative direction^38^. To minimize the effect of this attenuation, the AUC of the rate of change of the prestimulus baseline was used instead of the raw BOLD signal. This approach resulted in significant differences across many areas. As the brain regions of a participant stimulated by olfactory stimuli are influenced by inter-subject individual differences^39^ and the pleasantness of the odorous substance^40,41^, it is also important to look for several types of stimuli that will produce a more definite response.

Although the MD is the only olfactory pathway that does not respond, it thought to be associated with the link between olfaction and memory and is involved in olfactory stimulation identification and the association between olfactory stimulation and reward^42,43^. In this study, the olfactory stimulation of thiamine was not familiar to the marmosets; furthermore, it was not associated with rewards. Therefore, it is possible to consider that MD did not show a strong reactivity to this olfactory source. Additionally, the responses of Eth and Orbi are known to differ between the left and right sides and left-right differences are known from previous studies in humans^44,45^. This left-right difference was reproduced in the marmoset. This left-right difference in brain activity is known to be related to the comfort of olfactory stimulation^46^, and using various olfactory sources will broaden our understanding of the olfactory function of marmosets and primates.

No relationship was found between the olfactory pathway and the reaction times at each site. In particular, the peak in the orbitofrontal cortex, which is the endpoint of the olfactory pathway, was the fastest. Piri and Eth responded slower than Orbi. However, because olfactory response times are fast, on the order of milliseconds^47^, it is difficult to discuss response times from fMRI results, which have a temporal resolution of 1.5 s. In contrast, fMRI has superior spatial resolution and can capture responses throughout the brain. The lack of a predominant increase in responses in the thalamus in this study indicates that marmosets also have a pathway unrelated to the thalamus. Our fMRI data revealed the brain regions in marmosets that respond to olfaction. Therefore, it will be possible to conduct a more detailed study in the future by combining the results of EEG measurements with a high temporal resolution.

In this study, a significant response was observed in the OB, which is highly associated with PD^48,49^, and in the Eth, which is associated with AD^50^. Olfactory dysfunction is an early symptom of AD, and AD pathology in the entorhinal cortex, hippocampus, and temporal regions explains why patients with AD are unable to accurately identify odors^51^. Additionally, in patients with autosomal dominant AD, olfactory dysfunction has been shown to correlate with cognitive decline and the accumulation of tau in the entorhinal cortex^52^. In marmosets, there are PSEN1 mutant models^27^, a model of hereditary AD. The application of this device, which allows the observation of reactivity to olfactory stimulation, to model marmosets may allow the evaluation of whether olfactory dysfunction is useful as a biomarker of AD.

### Conclusions

We created a marmoset holder suitable for olfactory stimulation fMRI in an awake state. Its usefulness was evaluated in both mock and animal experiments. Ventilation was adequate, olfactory stimulation could be controlled on and off inside the MRI, and increased signal values in the brain regions were associated only with the olfactory pathway.

## Materials and Methods

### Animal Care and Management

This study was conducted in the animal facility of the Central Institute for Experimental Animals (CIEA) in Kawasaki, Japan. The animal experimental protocol was reviewed by the Institutional Animal Care and Use Committee and approved (approval no. 21129A) according to the Regulations for Animal Experiments of the CIEA, based on the Guidelines for the Proper Conduct of Animal Experiments by the Science Council of Japan (2006). Marmosets were kept in a cage in pairs or singly, depending on experimental and veterinary care reasons. The cage sizes were 409–820 × 610 × 728–1578 mm, and the cages were positioned facing each other to allow the animals to communicate visually and vocally. All the cages were equipped with sleeping areas, wooden perches, and hammocks. The animal rooms were conditioned at 26–28 °C and 40–60% humidity with a 12h:12h light/dark cycle. The animals tested negative for *Salmonella spp*., *Shigella spp*., and *Yersinia pseudotuberculosis* on annual fecal examinations.

### Visualizing airflow of olfactory stimulants

To visualize the degree of residual olfactory stimulants in a physical environment equivalent to that of an MRI experiment and to evaluate the usefulness of the suction, an MRI olfactory stimulation environment was established (Fig. 2 A, B, C). The simulation environment mimicked actual MRI to create an MRI olfactory stimulation environment consisting of an acrylic cylinder (Fig. 2-4) with an inner diameter of 74 mm and a total length of 650 mm that mimicked an MRI bore, a face mold made by molding a marmoset (Fig. 2-9), a marmoset holder (Fig. 2-3, 2-7), a receiver coil, (Fig. 2-13) and silicone supply and suction tubes (Fig. 2-11, -12) with an inner diameter of 4 mm. To visualize the gas flow, smoke (501, GASTEC) was used as a tracer particle source, and a laser light source (STS Corporation, GML-7010) with an irradiation angle of 70° was used. Images were recorded at 30 fps at a resolution of 1280 × 720 pixels. Transparent acrylic tubes imitating MRI bores were used to observe the tracer particles inside the tubes. The effect of the presence or absence of suction on the gas flow was visually investigated under conditions in which the supply and suction airflow rates were fixed at 0.3 L/min.

### Animal training

Prior to the fMRI experiment, all the animals were trained as described in a previous study^53^. Three marmosets (I7000, I814, and I817) were used for training and imaging (male, mean age was 6.7±1.9 years old; average body weight was 342.3±19.7 g). The marmoset in the holder was placed in a mock coil. Earphones were placed on top of the tube. The camera and light used to record the video were placed in the upper part of the coil. After the room was darkened and fMRI sounds were played, the training was started. The state of training was evaluated by the implementer for 1–3. The scoring criteria were as follows: 1. Did not move 2. Moved but did not interfere with the MRI 3. Movement that interferes with fMRI. The training began with a 15-min session. When the implementer scored the animals 1 or 2, the next training session was conducted for 30 min. If the training was performed for 45 min and the implementer scored the animal as 1 or 2, then fMRI was conducted. When the animal moved during the fMRI experiment, the training was repeated for 45 min. We recorded the animals during training using a video camera to confirm that the marmosets were awake. When the marmosets closed their eyelids, we confirmed their movement in response to environmental sounds. A liquid reward was provided after acclimatization. The operator carefully observed the animals before, during, and after acclimatization.

### Image acquisition

This study used a 7.0T Biospec 70/16 scanner system equipped with actively shielded gradients at a maximum strength of 700mT/m (Bruker biospin MRI GmbH; Ettlingen, Germany). The receiver coil was a 4-channel phased array surface coil, and the transmitter coil was a 72 mm quadrature transmit RF coil (Bruker BioSpin).

fMRI data were acquired under the following conditions: T2*-weighted functional images obtained using a single-shot gradient-recalled echo-planar imaging sequence for blood oxygenation level-dependent contrast (BOLD). The parameters were as follows: repetition time (TR)/echo time (TE): 1500/15 ms, field of view: 38.4× 38.4 mm, 21 axial slices, in-plane image Matrix 64 × 64, spatial resolution: 0.6 × 0.6 (mm), slice thickness 1 (mm), flip angle: 60°, repetitions 40 times, scan time: 60 s. A blip image was also taken to compensate for the fMRI data. Additionally, T2 Weighted Images (T2WI) were obtained to acquire anatomical images. The following parameters were obtained using rapid acquisition with a relaxation enhancement (RARE) sequence. TR/TE: 3000/30 ms, FOV: 40× 40 (xy plane) mm, 30 axial slices, image matrix 160 ×160, spatial resolution: 0.25 × 0.25 (mm), slice thickness: 1 (mm), RARE factor: 8, number of averages: 4, scan time: 3 min 12 s. The trained marmosets were subjected to fMRI. The subject was placed in the holder, restrained with a collar, and transported to the MRI room swiftly. When the animal was sufficiently calm, it was covered with inner and outer helmets. To observe vital signs, a respiration sensor was placed on the abdomen, and a pulse oximeter was placed on the tail. The supply and suction tubes were connected to a holder, which was placed inside an MRI machine. Imaging was performed for 60 s per trial, with olfactory stimulation turned off for the first 30 s and turned on for the next 30 s. The stimulus source was a thiamine propyl disulfide (Alinamin® injection 10 mg, Teva Takeda Pharma Ltd, Tokyo, Japan), which is used in human intravenous olfaction test^36^. The interval between the runs was 5 min. Suction was always applied during imaging and interval times. Each stimulation trial was repeated five times. The flow rate was 3 L/min for both the air supply and suction. To confirm that the signal declined and increased during stimulation, a 7-min scan was performed once. In this scan, stimuli were provided to the marmosets for 30 s and 6 min 30 s. A light source (Machida Endoscopy, Chiba, Japan) was placed in front of the animal’s face during the MRI. The light was necessary to keep the animals awake.

### Image analysis

After correcting the susceptibility distortion of the functional images using FSL TOPUP, slice and motion corrections were performed. Corrected functional images were normalized into the standard space using the parameters generated with T2WI normalization to the standard space using the ANTs suite package (Advanced Normalization Tools, http://stnava.github. io/ANTs/). Images were smoothed using an isotropic Gaussian kernel of 2.0 mm. Using Multi-image Analysis GUI (http://ric.uthscsa.edu/mango/download.html), time series data of the nine regions associated with olfaction (OB, AON, Piri, ACo, MD, EthR, EthL, OrbiR, and OrbiL) and four non-associated area (S1, Gu, V1, Aud) were extracted based on the MRI atlas.

### Statistical analysis

The signals in the ROI were normalized in each trial by the rate of change from a mean of 15 s immediately before stimulation. After normalization, to exclude the effect of baseline drift, the data were differentiated, and the area under the curve (AUC) between the derivative and 0 was calculated. Statistical analyses of the pre- and post-stimulus AUC values were performed using non-parametric paired t-tests (Wilcoxon signed-rank sum test). The threshold value for the acceptance of differences was 0.05.

## Acknowledgments

This research was partially supported by the Brain Mapping by Integrated Neurotechnologies for Disease Studies (Brain/MINDS), “Study of developing neurodegenerative model marmosets and establishment of novel reproductive methodology (JP19dm0207065)” from the Japan Agency for Medical Research and Development (AMED) and Grant-in-Aid for Early-Career Scientists “Effects of transient alcohol exposure to the fetus in early pregnancy on brain development.” JSPS KAKENHI, Grant Number JP20K16908 to TY. This work was partially supported by the Cooperative Research Project Program of the Joint Usage/Research Center at the Institute of Development, Aging, and Cancer, Tohoku University, and the Collaborative Research Project of the Institute of Fluid Science, Tohoku University.

## Data availability statement

Figure 3 data can be referred to Supplementary Data 1. Figure 4 data can be referred to Supplementary Data 2. Figures 5 and 6’s data can be referred to Supplementary Data 3.

## Contributions

T.Y., Y.I. and E.S. conceived and designed the study. T.Y., F.S., A.Y. and Y.I. developed the animal holder. A.Y., J.O., T.Y., Y.T., and Y.I. performed the mock circuit experiments. T.Y., F.S., M.K., I.T., and E.S. performed the MRI studies. T.Y., F.S., A.Y., T.I., Y.I. and E.S. discussed and interpreted the results. T.Y., F.S., A.Y., Y.I. and E.S. wrote the manuscript. All the authors commented on the manuscript.

