## Supplemental figure for "Development of a noninvasive olfactory stimulation fMRI system in marmosets": SupplementaryFig.pptx

### Slide 1
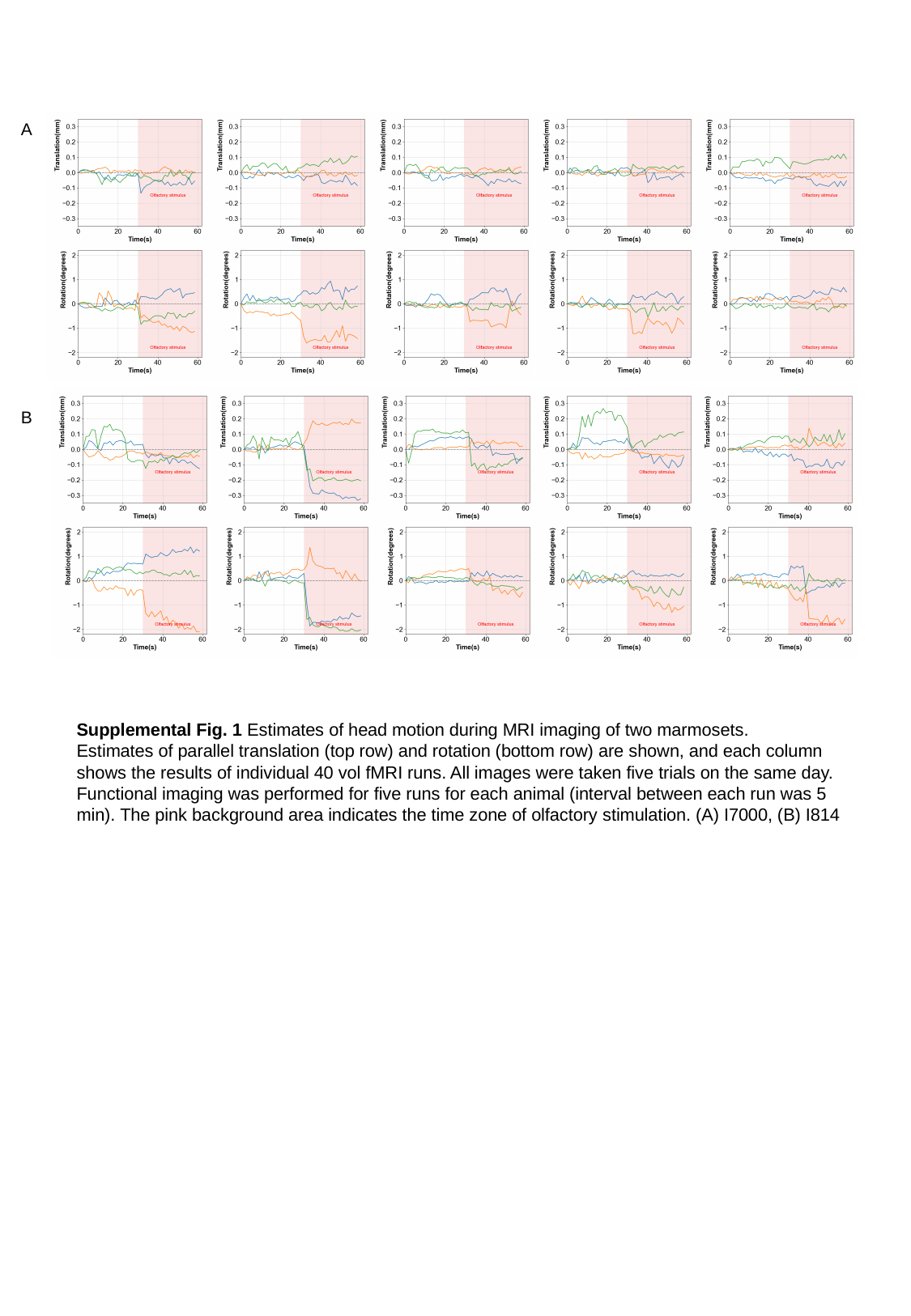

A
B
Supplemental Fig. 1 Estimates of head motion during MRI imaging of two marmosets.
Estimates of parallel translation (top row) and rotation (bottom row) are shown, and each column shows the results of individual 40 vol fMRI runs. All images were taken five trials on the same day. Functional imaging was performed for five runs for each animal (interval between each run was 5 min). The pink background area indicates the time zone of olfactory stimulation. (A) I7000, (B) I814

### Slide 2
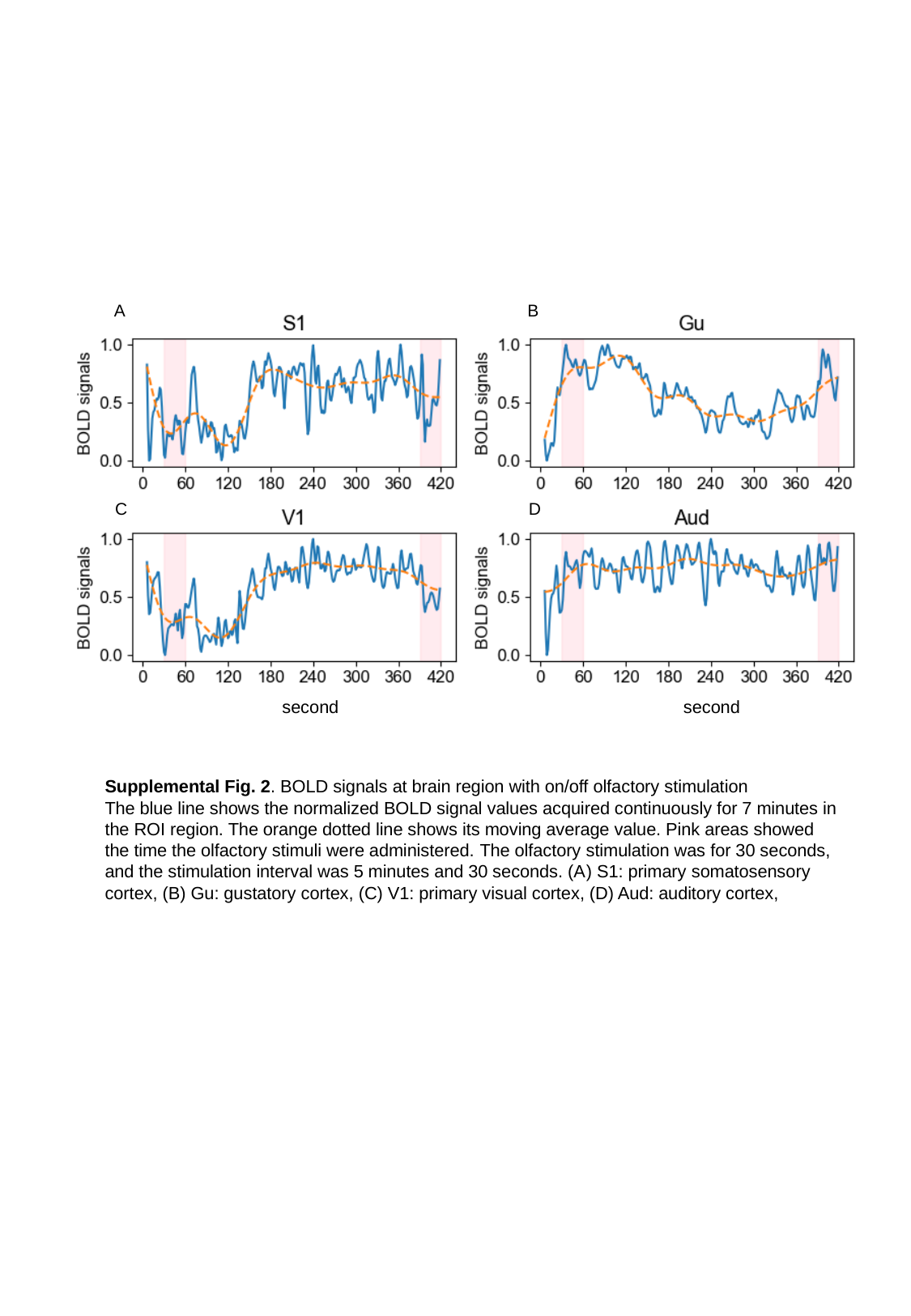

A
B
C
D
second
second
Supplemental Fig. 2. BOLD signals at brain region with on/off olfactory stimulation
The blue line shows the normalized BOLD signal values acquired continuously for 7 minutes in the ROI region. The orange dotted line shows its moving average value. Pink areas showed the time the olfactory stimuli were administered. The olfactory stimulation was for 30 seconds, and the stimulation interval was 5 minutes and 30 seconds. (A) S1: primary somatosensory cortex, (B) Gu: gustatory cortex, (C) V1: primary visual cortex, (D) Aud: auditory cortex,

### Slide 3
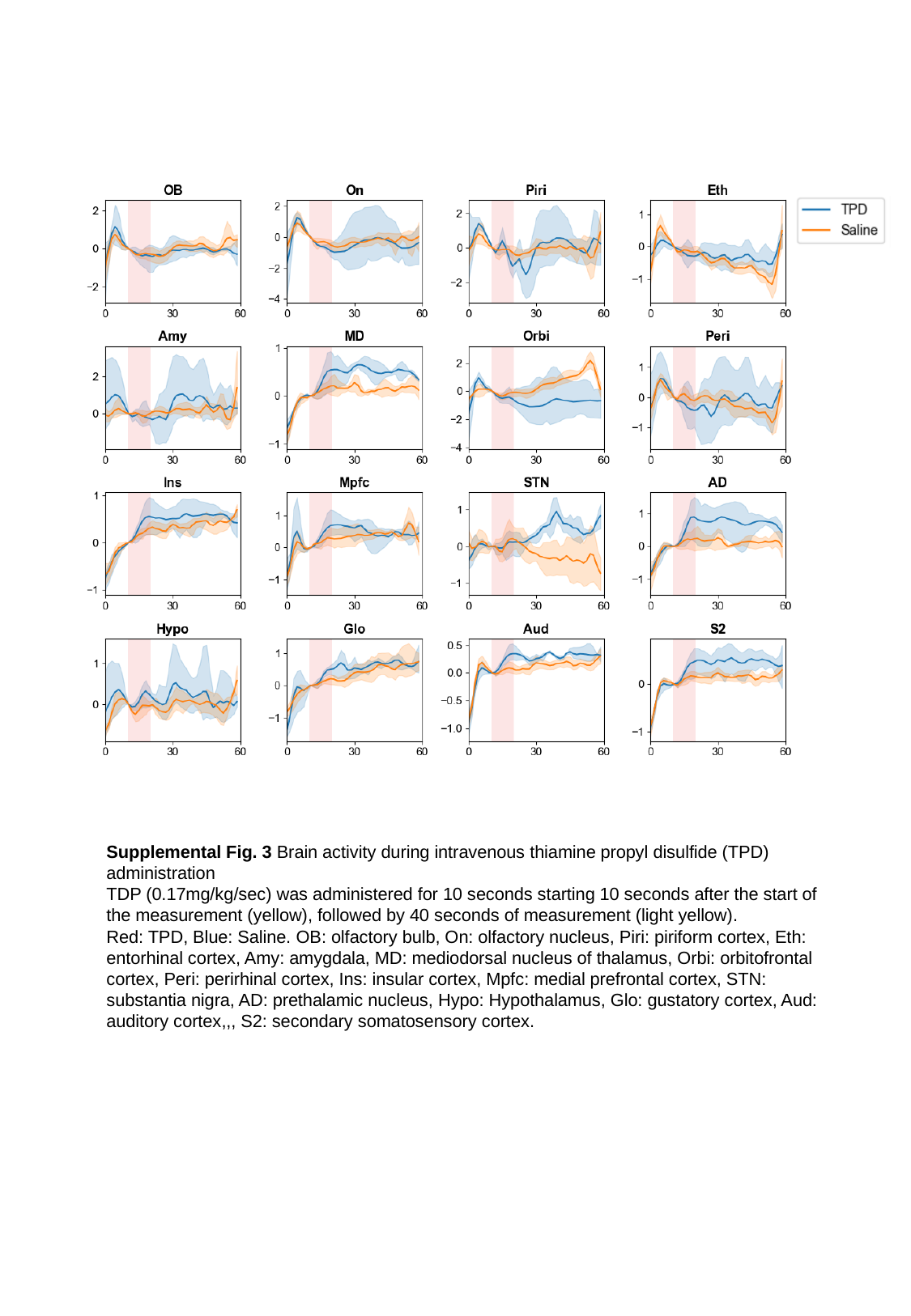

Supplemental Fig. 3 Brain activity during intravenous thiamine propyl disulfide (TPD) administration
TDP (0.17mg/kg/sec) was administered for 10 seconds starting 10 seconds after the start of the measurement (yellow), followed by 40 seconds of measurement (light yellow).
Red: TPD, Blue: Saline. OB: olfactory bulb, On: olfactory nucleus, Piri: piriform cortex, Eth: entorhinal cortex, Amy: amygdala, MD: mediodorsal nucleus of thalamus, Orbi: orbitofrontal cortex, Peri: perirhinal cortex, Ins: insular cortex, Mpfc: medial prefrontal cortex, STN: substantia nigra, AD: prethalamic nucleus, Hypo: Hypothalamus, Glo: gustatory cortex, Aud: auditory cortex,,, S2: secondary somatosensory cortex.
